# Lipid Droplet Targeting Drives Substrate Access during Very Long Chain Fatty Acid Activation by Fat1

**DOI:** 10.64898/2026.02.09.704383

**Authors:** Carolin Willner, Jennifer Sapia, Bianca M. Esch, Pia Erdbrügger, Lisa Kauffmann, Alicia Damm, Sergej Limar, Sebastian Eising, Stefan Walter, Abdou Rachid Thiam, Carolin Körner, Stefano Vanni, Florian Fröhlich

## Abstract

Fatty acids, the most abundant building blocks for lipid synthesis, require conjugation with Coenzyme A (CoA) for their incorporation into membrane lipids. Of those, very long chain fatty acids (VLCFAs), the precursors of yeast ceramides, are particularly hydrophobic. Yet it remains unresolved where and how the sole VLCFA-CoA synthetase Fat1 activates them. Here we show that lipid droplets (LDs) are both the site of free VLCFAs storage, as well as their place of activation. Free VLCFAs preferentially partition into LDs in molecular simulations and *in vitro* reconstitutions, whereas Fat1 targets the LD surface in cells via a N-terminal amphipathic helix. Structural predictions identify an essential hydrophobic cavity in Fat1 that connects the active site to the membrane, suggesting that VLCFAs directly shuttle from the LD core into the enzymatic pocket. Our model explains how the organelle targeting of a fatty acid activator evolved to match the biophysical properties and consequently the subcellular localization of its highly hydrophobic substrate.

## Introduction

Fatty acids (FAs) are the simplest class of lipids. Yet, they play multiple key roles in cell homeostasis, from acting as signaling molecules to serving as an energy reservoir for rapid cell growth via their role as building blocks for the most abundant membrane lipids, sphingolipids and glycerophospholipids^1^. FAs can vary in their length and also in their degree of saturation. The main FA species in the yeast *Saccharomyces cerevisiae* are the long chain fatty acids (LCFAs) harboring 16-20 carbon atoms and the very long chain fatty acids (VLCFAs) harboring 22-26 carbon atoms. The majority of yeast FAs are either saturated or mono-unsaturated. The LCFAs are substrates for glycerophospholipid biosynthesis enzymes as well as for the serine palmitoyl-transferase (SPT), the rate-limiting enzyme in sphingolipid biogenesis^2–4^. In contrast, the VLCFAs in yeast are mainly incorporated into ceramides and are also used in the biogenesis of GPI-anchored proteins^5,6^.

FAs are either taken up by cells from the extracellular space or can be directly synthesized by cells. The process of FA uptake is rather unclear. FAs are either integrated in the plasma membrane and enter the cell via passive diffusion or via active transport mechanisms. In mammalian cells, the CD36 family of proteins are thought to be involved in FA uptake^7^. In yeast, a process called vectorial acylation, that depends on the FA CoA-synthetase family of proteins^8^, has been proposed as the main mode of FA uptake. It is believed that FAs enter the cell in their non-activated form, generally described as free FAs. Free FAs need to be activated by the addition of a Coenzyme A (CoA) molecule. This reaction is catalyzed by the FA CoA synthetase protein family in an ATP-dependent reaction forming a high energy thioester between the carboxyl group of the FA and the CoA molecule^9–11^.

In parallel, FAs can also be synthesized by cells. The rate-limiting step in FA biosynthesis is catalyzed by the acetyl-CoA carboxylase (Acc1), which catalyzes the carboxylation of cytosolic acetyl-CoA to form malonyl-CoA^12,13^. Malonyl-CoA and acetyl-CoA are the substrates of the fatty acid synthase (FAS) that extends growing FAs in an iterative process by two carbons at a time. The final product, typically palmitic acid, is released from the acyl-carrier protein as a free FA that has to be activated^14–16^. In a similar process, LCFAs can be further elongated up to 26 carbons by the VLCFA elongation complex at the ER. In contrast to the FA synthase, this process releases CoA-activated VLCFAs^17^.

In addition to the *de novo* biosynthesis pathway, free FAs are also generated by the action of various lipases. Triacylglycerol lipases generate free FAs at lipid droplets (LDs), the lipid storage organelles of cells^18,19^. Free VLCFAs are generated mainly by the action of ceramidases. While mammals harbor acidic, neutral and alkaline ceramidases^20,21^, yeast only expresses the two alkaline ceramidases Ypc1 and Ydc1^22^. Very little is known about the biophysical behavior and the intracellular localization of free VLCFAs. In yeast, the CoA-dependent activation relies exclusively on the VLCFA acyl-CoA synthetase Fat1^23^. This enzyme has been proposed to be localized at the plasma membrane, peroxisomes and LDs^24–27^. However, its membrane targeting mechanism, its exact intracellular localization and where and how it activates free VLCFAs are largely unknown.

Here, we investigate the behavior of free VLCFAs in a membrane environment. Computational and biophysical data suggest that free VLCFAs preferentially partition into LDs. We demonstrate that Fat1 harbors an N-terminal amphipathic helix with a high hydrophobic moment that binds to both the LD monolayer and the endoplasmic reticulum (ER) with a strong preference for LDs. Based on structural predictions, we show that Fat1 harbors a hydrophobic cavity that spans the entire protein from the active site to the membrane-binding amphipathic helix. Taken together, our data support a model in which free VLCFAs, upon release from ceramides, spontaneously partition into LDs and enter the active site of Fat1 from the core of the LD. This suggests that the unusual LD targeting motif of Fat1 co-evolved with the biophysical properties of free VLCFAs.

## Results

### The biophysical properties of very long chain fatty acids promote their enrichment into lipid droplets

In *S. cerevisiae*, free VLCFAs are produced by the ceramidases Ypc1 and Ydc1 via the hydrolysis of ceramides to generate a sphingoid base and free FA. In contrast to mammalian cells, the two yeast ceramidases belong to the family of alkaline ceramidases, and both Ydc1 and Ypc1 have been suggested as ER resident proteins^22^. However, these studies were performed under strong overexpression conditions and without comparison to an ER marker^22,28,29^. To confirm the co-localization of Ydc1 with the ER and to understand where in yeast cells free VLCFAs are generated, we tagged Ydc1 N-terminally with an mNeon tag under the control of the Pho5 promoter and analyzed its localization in correspondence to DsRed-HDEL, a common ER marker. Our analysis revealed high co-localization of mNeon-Ydc1 with DsRed-HDEL, strongly suggesting that free VLCFAs are formed at the ER (**Fig. 1A**). Once generated, VLCFAs are incorporated into membrane and storage lipids or transported to peroxisomes for their degradation via β-oxidation^30,31^.

**Figure 1.**
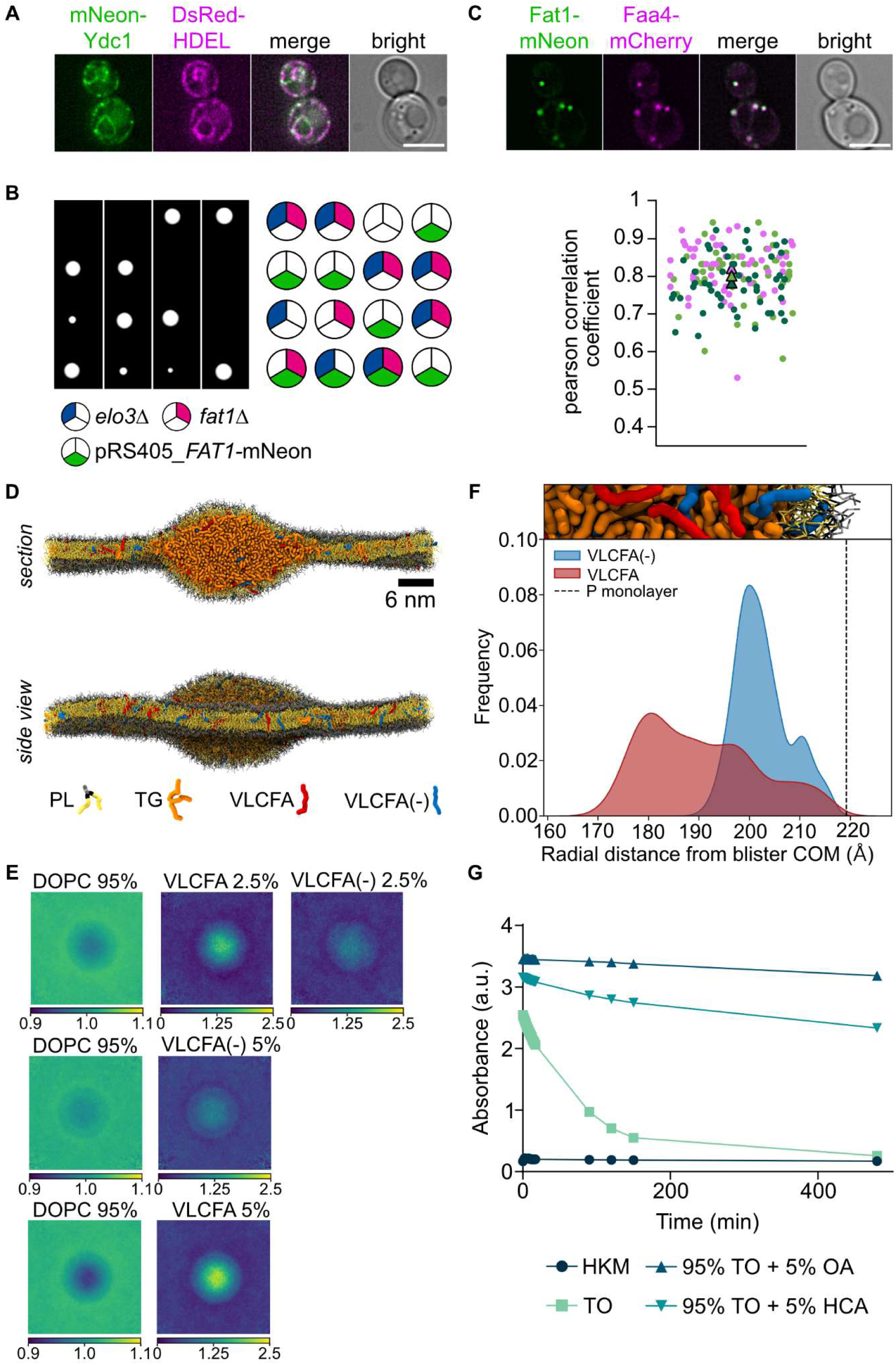
**A)** Co-localization of mNeon-Ydc1 and the ER marked with DsRed-HDEL in WT cells. scale bar = 5 µm. **B)** Tetrad analysis of fat1Δ pRS405-FAT1-mNeon cells (pink and green, respectively) mutants crossed with elo3Δ mutants (blue). **C)** Co-localization of Fat1-mNeon with the LD marker Faa4-mCherry. Representative confocal midsections are shown (upper panel). Scale bar = 5 µm. Quantification of the co-localization of Fat1-mNeon and Faa4-mCherry using the Pearson correlation coefficient (lower panel). Three independent experiments are shown (dark green, green, pink). Each dot represents a single analyzed cell. Each triangle represents the mean from each experiment. **D**) Top, side and section views of CG-MD simulation starting set up for a nascent LD-like ternary system, comprising a TG oil core (orange) covered by PLs (head: gray; tails: yellow). The system is enriched with 2.5% neutral (red) and 2.5% anionic (blue) VLCFAs. Solvent is not shown for clarity. **E)** Enrichment-depletion 2D maps showing the accumulation of both neutral and charged VLCFA molecules at the LD lens. **F)** Radial distribution distance of VLCFA in both neutral and anionic form from the triolein (TO) blister COM. **G)** Stability measurements (λ = 340nm) of triolein (TO) droplets with and without FAs over time. Oleic acid (OA) stabilizes a TO droplet better than hexacosanoic acid (HCA). TO droplets are instable. HKM = buffer control.

However, all these processes require the activation of the free FA. In yeast, Fat1 is the sole acyl-CoA synthetase that specifically activates C20-C26 fatty acids^32^. Previous studies have described Fat1 as a transmembrane protein localized at the ER, the plasma membrane as well as LDs^24,26^. To exactly characterize the intracellular localization of Fat1, we generated a C-terminally mNeon tagged Fat1 construct. We first confirmed functionality of the tagged protein by its ability to rescue the synthetic phenotype of a *FAT1 ELO3* double deletion^33^. The double deletion of *ELO3* and *FAT1* failed to produce any viable progeny in tetrad dissections **(Fig. 1B)**. This phenotype was rescued by expressing Fat1-mNeon from a plasmid under its endogenous promotor **(Fig. 1B)**. Expressing wildtype (WT) Fat1 from a plasmid yielded similar results (**Sup. Fig. 1A**). We next analyzed the localization of Fat1-mNeon in yeast cells. We observed a dot-like localization and failed to observe the typical ER or plasma membrane signal. To test whether the observed structures were LDs, we analyzed the localization in respect to the LD marker Faa4-mCherry. Fat1-mNeon showed high co-localization to Faa4-mCherry with an average Pearson correlation coefficient of 80% **(Fig. 1C)**. Thus, our data show that Fat1 is a LD protein. While previous studies proposed a role for Fat1 in the uptake of free VLCFAs from the medium^8^, its strict localization to droplets challenges this hypothesis.

If Fat1 is not involved in the uptake of VLCFAs, our results raise the question if free VLCFAs partition to LDs, where they would be activated by Fat1. With 22-26 carbon atoms, the free VLCFAs that are generated by ER ceramidases are very hydrophobic molecules that harbor only a small polar carboxyl group, which protonation depends on the length of the acyl chain^34^. Since direct imaging of VLCFAs in cells is extremely challenging due to their highly dynamic chemical nature, we turned to molecular dynamics (MD) simulations and an interfacial stability assay to assess partitioning tendencies. To this end, we modeled C24:0 FAs in both neutral and charged forms and analyzed partitioning of VLCFAs in a MD system containing both a lipid bilayer and a nascent LD blister **(Fig. 1D)**. Coarse grain (CG) simulations reveal that protonated VLCFAs almost exclusively partition to the neutral core of nascent LD blisters, strongly favoring the oil phase (**Fig. 1E**). Charged C24:0 FAs accumulate in the region surrounding the TG blister, whereas the bilayer regions were notably depleted of FAs (**Fig. 1E-F, Sup. Fig. 1C**). Analysis of the radial distribution and solvent-accessible surface area (SASA) revealed that both VLCFA species (neutral and anionic) populate regions within the droplet core as well as near the phospholipid monolayer. The anionic form shows a relatively increased enrichment toward the droplet surface, likely indicating a possible preference for localization at the interface between the neutral lipid core and the phospholipid shell (**Fig. 1F**). In summary, this suggests that ER-generated VLCFAs tend to preferentially partition into LDs compared to the ER membrane.

To confirm our MD simulations, we analyzed the capacity of VLCFAs to be recruited to the oil/water interface of triglyceride oil-in-buffer emulsion droplets. The coverage of such interface by surfactant molecules would improve the stability of the emulsion which we measured by recording the absorbance at 340 nm of the emulsion as a direct readout for droplet stability. While a TG-only droplet becomes unstable over time, the addition of 5% C26:0 hexacosanoic acid stabilized the emulsion, suggesting some surfactant-like properties of the FA (**Fig. 1G**). However, the effect is not as strong as for oleic acid (OA), which appears to be a better surfactant (**Fig. 1G**). Together, our results suggest that ceramidase-derived free VLCFAs form at the ER and are predicted to spontaneously partition into LDs. While neutral VLCFAs are found preferentially in the core of LDs, deprotonated VLCFAs have surfactant-like properties and will spontaneously appear at the monolayer of LDs.

### Fat1 targets lipid droplets via a hydrophobic amphipathic helix

We next investigated how Fat1 is targeted to LDs. To explore how Fat1 interacts with membranes, we employed CG-MD simulations. In the absence of an experimentally resolved 3D structure for the yeast Fat1 protein, we utilized the Alphafold2 (AF2)^35,36^ model **(Fig. 2A)**, which was predicted with high overall accuracy (pLDDT score > 90). However, regions such as the N-terminal helix and several loops were predicted with lower confidence, likely reflecting their inherent flexibility, as shown for other membrane targeting proteins of similar fold^37^.

**Figure 2.**
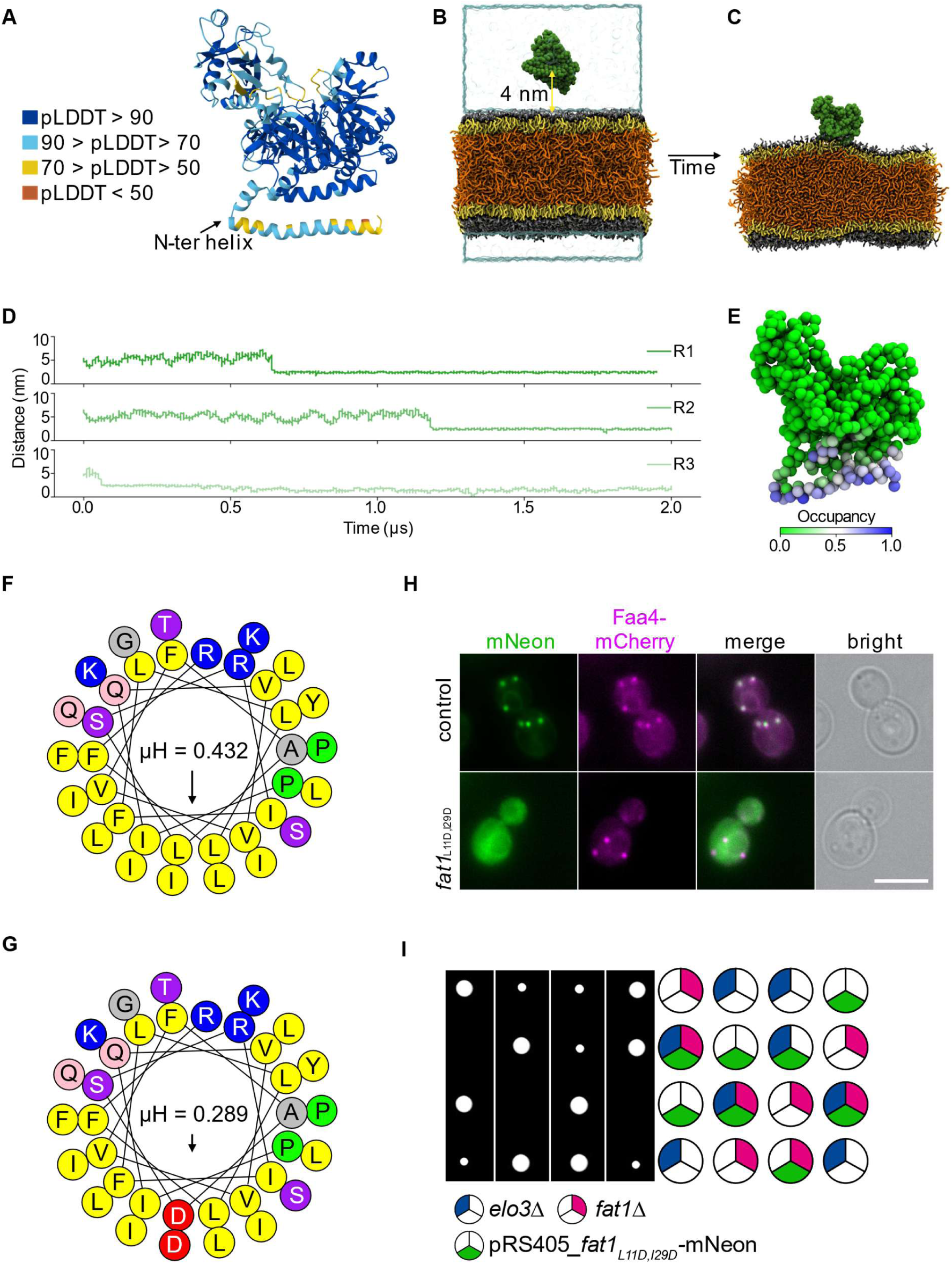
**A)** AF2 model of yeast Fat1. Color code based on pLDDT score. **B-C)** CG-MD simulation setup **(B)** showing Fat1 in solution in the presence of a trilayer. (PL headgroups: grey; PL tails: yellow; TG: orange) and subsequently bound to the model-LD monolayer **(C)**. **D)** Time trace of minimum distance values all replicates, interacting via its N-terminal helix. **E)** Occupancy map showing the regions in contact with the bilayer (blue). **F)** and **G)** Helical wheel representation of the first 3-37 amino acids. Hydrophobic residues are shown in yellow and the hydrophobic moment (µH) is depicted as an arrow, with the corresponding value shown above. **(F)** WT sequence and **(G)** *fat1*_L11D,I29D_ mutant, in which the hydrophobic residues leucine 11 and isoleucine 29 were replaced by charged aspartates (red). The projection was generated by the Heliquest software (http://heliquest.ipmc.cnrs.fr). **H)** Co-localization of Fat1-mNeon or *fat1*_L11D,I29D_-mNeon and LDs marked with Faa4-mCherry. Scale bar = 5 µm. **I)** Tetrad analysis of *fat1*Δ pRS405-*fat1*_L11D,I29D_-mNeon cells (pink and green respectively) mutants crossed with *elo3*Δ (blue).

As Fat1 binds to LDs, we simulated Fat1 in the presence of a trilayer system resembling an LD-like environment, consisting of a TG core sandwiched between two phospholipid monolayers **(Fig. 2B)**^38–42^. Initially localized in solution with a random orientation, approximately 4 nm from the trilayer **(Fig. 2B)**, Fat1 was consistently observed to interact with the trilayer via its N-terminal α-helix **(Fig. 2C**) in all MD replicates **(Fig. 2D)**. Quantitative analyses confirm a strong interaction at the N-terminal domain, supporting its role in membrane association **(Fig. 2E)**.

While this region was previously predicted to be transmembrane^24^, both the AF model (**Fig. 2A**) as well as our simulations (**Fig. 2B-E**) suggest that the N-terminal part forms an amphipathic helix. Heliquest also predicts the first α-helix of Fat1 to form an amphipathic helix **(Fig. 2F)**^43^. Helical wheel plots of the AAs 3-37^43^ reveal a strong amphipathic character of the long α-helix with a hydrophobic moment of µH=0.432, high hydrophobicity H=0.962 and a net charge of +4 **(Fig. 2F)**, which are characteristic parameters for LD binding amphipathic helices^44^. To disrupt the amphipathic character of the N-terminal Fat1 helix, we changed two hydrophobic residues, leucine 11 and isoleucine 29, to aspartic acid (L11D, I29D). These mutations reduce the hydrophobic moment of the Fat1 N-terminal helix to µH=0.289 **(Fig. 2G)**. We analyzed the localization of the mNeon-tagged *fat1_L11D,I29D_* construct in comparison to LDs marked with Faa4-mCherry. As expected for an amphipathic helix protein, changing the hydrophobic moment of the Fat1 N-terminal helix caused the protein to localize to the cytoplasm and not to LDs as observed for the WT protein **(Fig. 2H)**. Next, we tested the functionality of the *Fat1_L11D,I29D_*-mNeon construct by expressing it in the background of the *fat1Δ elo3Δ* double mutant. Expression of the construct was not sufficient to rescue the synthetic lethality **(Fig. 2I**), suggesting that the *fat1_L11D,I29D_*-mNeon protein is not functional when located in the cytoplasm. In addition, the N-terminal region of Fat1 fused to mNeon was sufficient to target LDs marked with Faa4-mCherry **(Sup. Fig. 1B**, lower panel), while deletion of the N-terminal domain resulted in a cytosolic localization **(Sup. Fig. 1B**, middle panel). In summary, our results suggest that the N-terminal domain of Fat1 forms an amphipathic helix that allows targeting of the protein to LDs.

### The amphipathic helix of Fat1 targets both lipid droplets and the endoplasmic reticulum

Many LD proteins with amphipathic helices target LDs from the cytoplasm, suggesting that the helix folds in the presence of a phospholipid monolayer^45,46^. However, the strong hydrophobicity of the Fat1 N-terminal helix suggests low solubility in the cytoplasm. We therefore investigated the localization of Fat1 in cells lacking

LDs. We analyzed the localization of Fat1-mNeon in a LD deficient strain that lacks the two diacylglycerol (DAG) acyl-transferase genes *DGA1* and *LRO1* and the two acyl-CoA sterol acyltransferase genes *ARE1* and *ARE2* (*Δ4*)^47^ and compared it to a WT strain. In the WT strain, Fat1 localized in dot-like structures adjacent to the ER marked with DsRed-HDEL (**Fig. 3A**, upper panels). In the *Δ4* strain, Fat1 localized to the ER based on co-localization with the DsRed-HDEL ER marker (**Fig. 3A**, lower panels) suggesting Fat1 targets the ER in the absence of LDs. To confirm our results, we repeated our CG-MD simulations of Fat1 in a bilayer system. We again initially localized Fat1 in solution with a random orientation, approximately 6 nm from the bilayer. Over time, Fat1 interacted with a simple DOPC bilayer, again via its N-terminal amphipathic α-helix **(Fig. 3B)**. The localization of Fat1 in LD harboring cells suggests a preference of the amphipathic helix for a monolayer over a bilayer. To confirm our results *in silico*, we simulated a ternary system where both monolayer and bilayer membranes coexist **(Fig. 3C)**^48^. This setup included a TG lens encapsulated between two monolayers, formed from the unzipping of a bilayer. Fat1 was initially positioned in solution, starting from 9 different initial localizations (**Supp. Fig. 2B**) and the time evolution of the protein was analyzed, focusing on bilayer *vs* lens localization (**Fig. 3C-E**). The results from the lateral distance between the protein and the TO core indicates that the protein progressively approaches the oil core during the simulations (**Fig. 3D**). Furthermore, center of mass (COM) tracing and *in silico* colocalization analysis show that the protein has a clear preference for binding to the monolayer region, where in most cases it remained stably associated with the TG blister **(Fig. 3E, Sup. Fig. 1E)**. Overall, of the 18 replicas, 10 of them end up in the LD lens after 1 μs of simulation time, despite only 2 of them starting directly on top of the LD lens (**Sup. Fig. 1D**). Moreover, the binding interface observed in this system matched that of previous simulations **(Fig. 3C, inset)**, further confirming the critical role of the N-terminal helix in facilitating the protein’s attachment to membranes.

**Figure 3.**
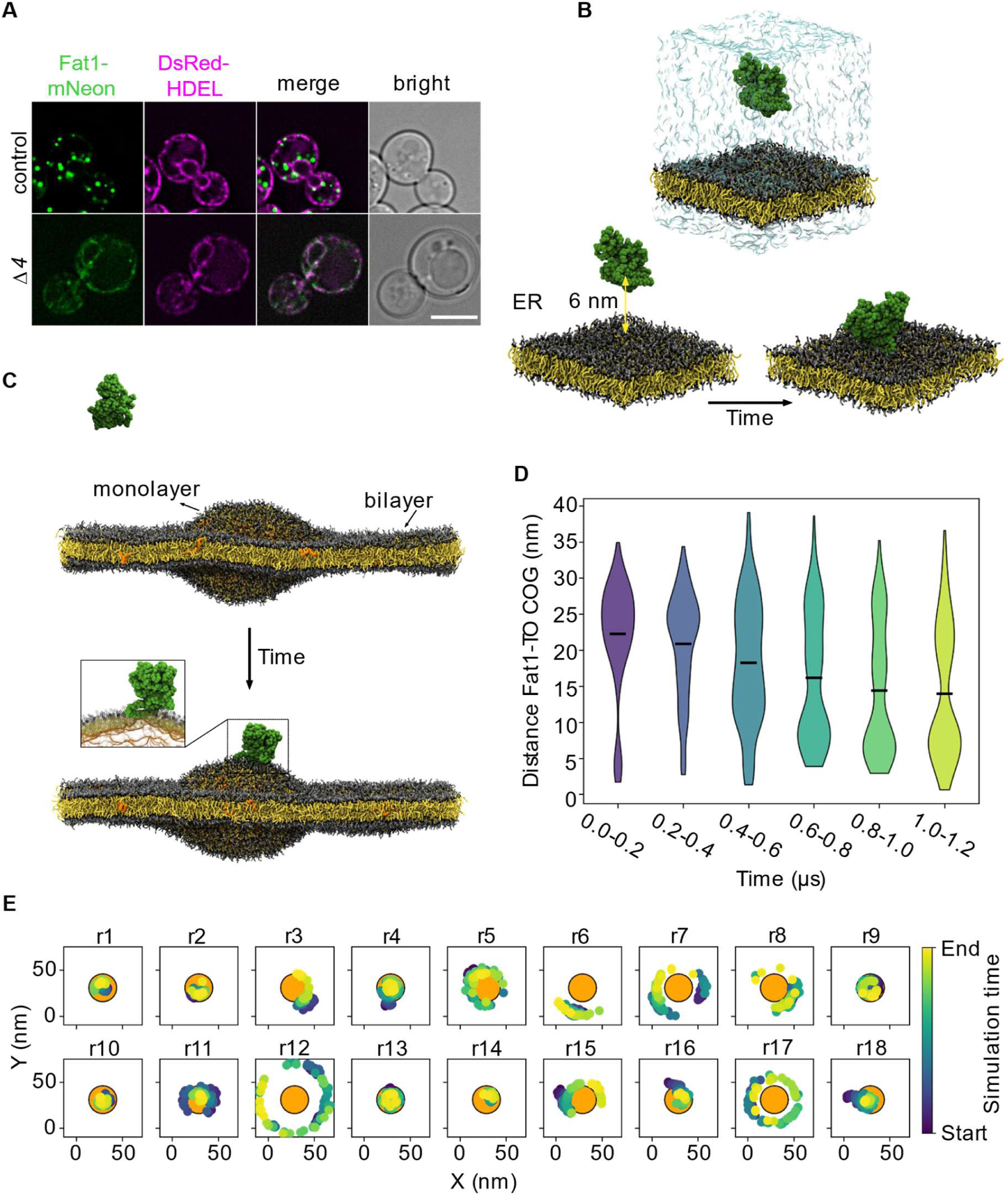
**A)** Co-localization of Fat1-mNeon and the ER marker DsRed-HDEL in WT cells (upper panel) and lipid droplet deficient cells (*Δ4*; *dga1*Δ *lro1*Δ *are1*Δ *are2*Δ; lower panels). Scale bar = 5 µm. **B)** CG-MD simulations setup and representative mechanism of yeast Fat1 (green) binding to a DOPC monolayer (PL headgroups: grey; PL tails: yellow). **C-E)** Fat1 binds preferentially to monolayer regions in a nascent LD model. **C)** Representative snapshots of Fat1 binding. The inset shows how Fat1 binds to monolayer through its N-terminal domain. **D)** Lateral distance between protein and TO-core centers of geometry (COG) over time. Violin plots show the distribution of x/y-plane distances between the protein and the TO-core across all 18 simulation replicas, divided into six equal time segments of 0.2 μs each. The black lines indicate the mean distance within each interval. The color code indicates the simulation time (violet=start, yellow=end). **E)** Fat1 center-of-mass (COM) trajectories in the x/y plane aligned to the TO blister COM (orange circle). Each panel shows the motion of an independent replica, with color indicating the simulation time (violet=start, yellow=end). Two sets of replicas were run for each of the nine initial positions (see **Sup. Fig. 1D**).

### Fat1 positioning on membranes is important for its function

Our results demonstrate that Fat1 is targeted to LDs or the ER via its amphipathic N-terminal helix. This targeting mechanism brings it together with its substrate, the free VLCFA. Similar to Fat1, free VLCFAs have a strong preference for LDs over the ER, suggesting that the biophysical properties of the amphipathic helix are adapted to those of the FAs. But how do free VLCFAs leave LDs to enter the catalytic cavity of Fat1? To answer this question, we again utilized the AF2 model of Fat1 to predict hydrophobic cavities that would be sufficient to harbor a VLCFA. The analysis revealed a tunnel extending from the protein surface, situated just behind the predicted position of the N-terminal helix, leading deep into the predicted^49^ putative active site with highly conserved CoA- and ATP/AMP binding regions **(Fig. 4A, Sup. Fig. 1F)**.

**Figure 4.**
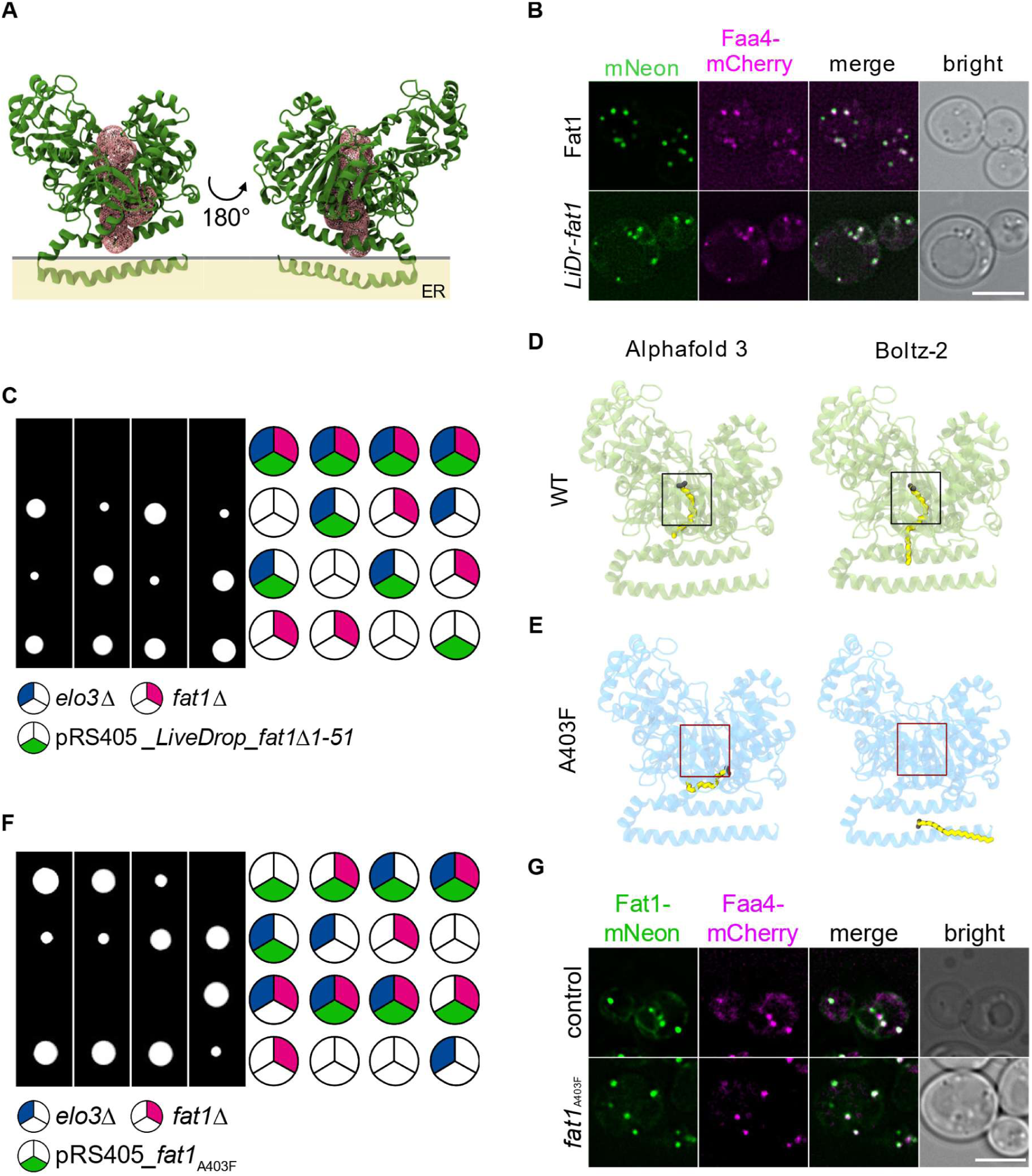
**A)** Cavity detection analysis shows the presence of a tunnel which extends from the Fat1-membrane binding interface deep into the protein core. **B)** Co-localization of Fat1-mNeon or *LiveDrop*-*fat1*-mNeon and LDs marked with Faa4-mCherry. Scale bar = 5 µm. **C)** Tetrad analysis of *fat1*Δ pRS405-*LiveDrop*-*fat1Δ1-51* cells (pink and green, respectively) mutants crossed with *elo3*Δ (blue). **D)** AF3 ^50^ and Boltz-2 ^51^ predictions of WT-Fat1 bound to a VLCFA (yellow licorice; oleic acid and lignoceric acid, respectively). **E)** AF3 ^50^ and Boltz-2 ^51^ predictions of A403-Fat1 show the incapability of the substrate VLCFA (yellow licorice; oleic acid and lignoceric acid, respectively) to bind into the predicted binding cavity. **F)** Tetrad analysis of *fat1*Δ pRS405-*fat1*_A403F_ cells (pink and green respectively) crossed with *elo3*Δ mutants (blue). **G)** Co-localization of Fat1-mNeon or *fat1*_A403F_-mNeon and lipid droplets marked with Faa4-mCherry. Scale bar = 5 µm.

Furthermore, the binding mode adopted by the protein when associated to a membrane appears to facilitate the opening of the detected cavity towards the membrane, suggesting a potential role in substrate recruitment. In line with these predictions, exchanging the first 51 amino acids of Fat1 with the *LiveDrop* LD targeting sequence^52–54^ allows the recruitment of the protein to LDs **(Fig. 4B)**, but renders the protein inactive. Expression of the chimera yielded no viable progenitors in tetrad dissections of *ELO3 FAT1* double deletions **(Fig. 4C)**. To further verify whether this cavity might indeed act as a binding pocket for VLCFAs, we modeled Fat1 in the presence of VLCFAs using AI-tools such as Alphafold3^50^ and Boltz-2^51^, which allow accurate predictions of how proteins interact with small molecule ligands. These models suggest that VLCFAs bind to the cavity we previously identified (**Fig. 4A**) with their carboxyl groups pointing towards the cytosolic domain **(Fig. 4D)**.

To test this model, we generated a tunnel mutant in which we exchanged A403 to a bulky phenylalanine (F) to create a steric hindrance and prevent the entry of the VLCFA substrate. In AI models of this tunnel mutant (A403F), the deep binding of the VLCFA was indeed hindered, with the ligand fully excluded from the binding cavity **(Fig. 4E)**, suggesting that the point mutation near the entrance of the active site might obstruct penetration of the FA into the cavity. In line with this, expressing the *fat1_A403F_* mutant did not rescue the synthetic lethality of the *FAT1 ELO3* double mutant (**Fig. 4F**), while the protein still localized to LDs (**Fig. 4G**). Importantly, neither tagging Fat1 with mNeon, nor changing alanine 403 to phenylalanine affected the expression levels of the protein (**Sup. Fig. 2 A,B and Sup. Tab. 4)**. Taken together, these observations point to the identification of a binding pocket inside Fat1, which opens up towards the membrane and is able to accommodate the VLCFA substrate.

To directly compare the activity of the WT Fat1 protein and the A403F mutant, we purified the two FLAG-tagged proteins after overexpression from the *GAL1* promotor in the presence of detergent^3^. Both purifications yielded similar amounts of proteins, suggesting that the A403F tunnel mutant is indeed stable in cells (**Fig. 5A**). We next established an *in vitro* activity assay based on the consumption of ATP and the detection of AMP release during the reaction. The molecular mechanism of fatty acid activation requires binding of ATP, binding of the fatty acid, formation of an acyl-AMP, release of the pyrophosphate, release of AMP and finally the formation of the thio-ester between CoA and the fatty acid^32^ **(Fig. 5B)**. In the assay, we mixed ATP, free CoA and α-cyclodextrin solubilized C26 FAs with the purified proteins. When we compared wildtype Fat1 and *fat1_A403F_* we observed around 7 to 9-fold increased activity of the WT protein over the mutant (**Fig. 5C**). However, this increase was directly depending on the amount of Fat1 added and the calculated amount of produced AMP suggests a pre-loading of the purified proteins with C26 FAs. In line with this, we detected similar activities when we did not add any C26 FAs as a substrate for the assay. Since the assay was performed in solution in presence of detergent this suggests that the generated C26-CoA cannot leave the binding pocket in the absence of membranes or proteins binding to C26-CoA. Alternatively, C26 FAs may not be sufficiently soluble, even in the presence of α-cyclodextrin, and thus, no substrate is available for the *in vitro* assay. Interestingly, the assay also produced AMP in the absence of additionally added CoA, suggesting that the purified proteins are bound to both, a C26 FA and CoA. However, this is not in line with the proposed Bi Uni Uni Bi Ping-Pong mechanism for acyl-CoA synthetases, where CoA is only supposed to bind after the AMPylation of the FA substrate^55^. In our hands, only the absence of ATP in the reaction led to no detectable signal. To test if the solubility of the FA substrate is a problem, we performed the assay with palmitate (C16) instead of C26 FAs as a substrate. In this case, we detected further substrate turnover suggesting that the FA can enter the active site and the CoA-activated product can leave as a soluble molecule. Here, the activity of the WT protein was significantly higher than the A403F mutant (**Fig. 5D**) suggesting that even a shorter C16 long chain fatty acid poorly partitions into the lipid binding tunnel of the mutant protein compared to the WT protein. In summary, the biochemical characterization of the WT Fat1 protein and the A403F mutant support the model of a long hydrophobic cavity necessary for the accommodation of the substrate in the enzyme.

**Figure 5.**
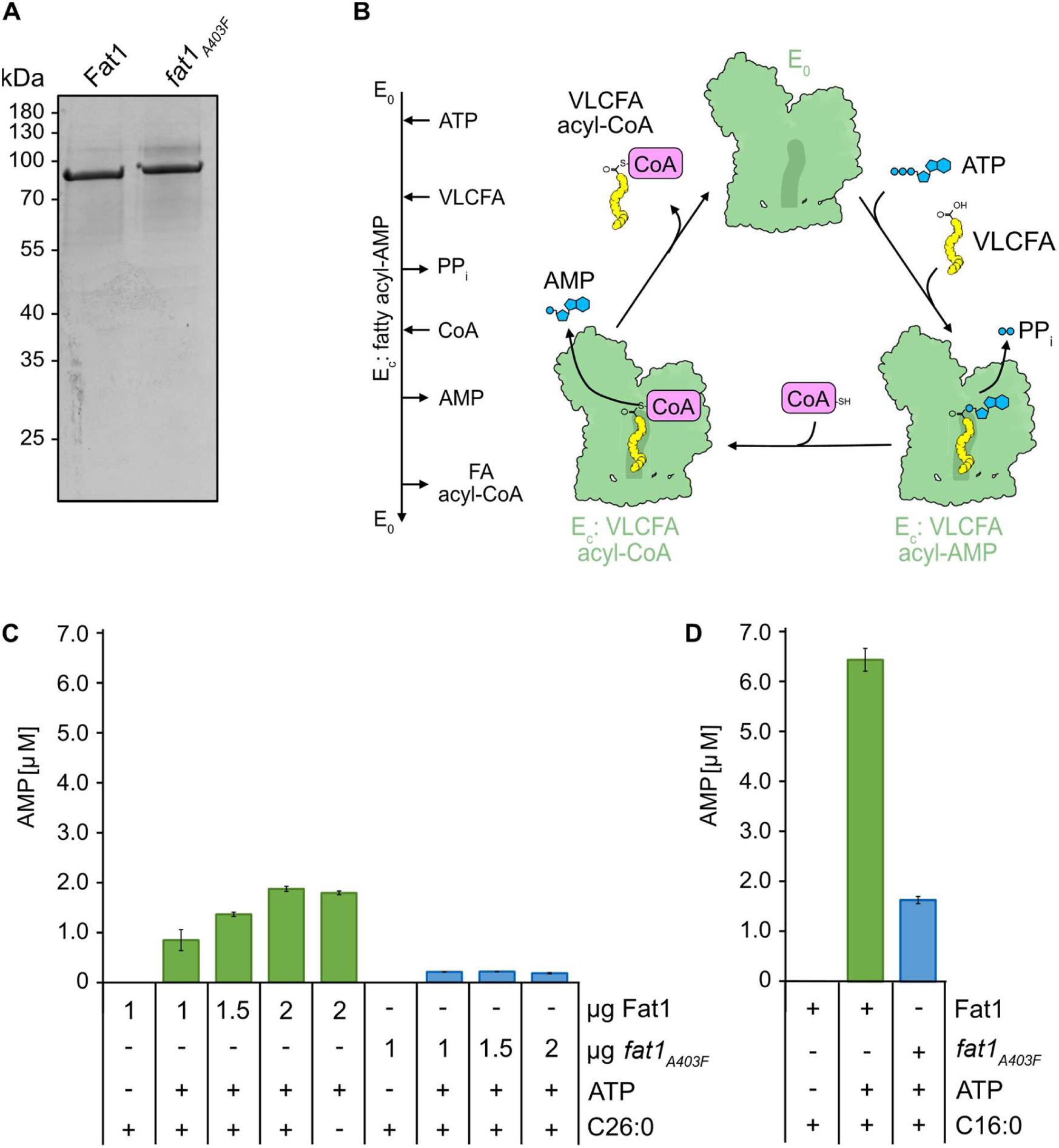
**A)** Coomassie-blue stained SDS-PAGE gel of purified Fat1-3xFLAG and *fat1*_A403F_-3xFLAG (n=1). **B)** Schematic overview of the proposed molecular mechanism of FA acid activation catalyzed by Fat1. The reaction is initiated by the formation of the acyl-adenylate intermediate via the binding of ATP and its VLCFA substrate resulting in the release of pyrophosphate (PP_i_). In the next step, AMP is exchanged for CoA leading to the production of the VLCFA acyl-CoA. **C)** AMP levels (µM) were measured from reactions containing increasing amounts of purified Fat1 or the tunnel mutant *fat1*_A403F_ in the presence or absence of ATP and C26:0. Wild-type Fat1 produces AMP in a concentration-dependent manner (green bars), even in the absence of the C26:0 FA substrate. The A403F tunnel mutant shows minimal activity (blue bars). Error bars represent standard deviations from three technical replicates (n=3). **D)** AMP levels (µM) were measured from reactions containing 1 µg of purified Fat1 or of the tunnel mutant *fat1*_A403F_ in the presence or absence of ATP and C16:0. Wild-type Fat1 produced significantly higher levels of AMP compared to the A403F tunnel mutant. Error bars represent standard deviations from three technical replicates (n=3).

## Discussion

VLCFAs are important building blocks of ceramides and GPI-anchored proteins in yeast. They are synthesized in their active form via the VLCFA elongation cycle. Alternatively, free VLCFAs are generated in the ER by the activity of the yeast ceramidases Ypc1 and Ydc1. This pool of free VLCFAs has to be activated by the VLCFA-CoA synthetase Fat1. Here, we used MD simulations and *in vitro* experiments to show that highly hydrophobic free VLCFAs preferentially partition to LDs due to their physicochemical properties **(Fig. 1)**. The pool of free VLCFAs generated by the ER-localized ceramidases in yeast is perfectly positioned for the partitioning since LDs in yeast pre-dominantly stay connected to the ER^56,57^. Free VLCFAs likely carry a charge on their deprotonated carboxyl group which gives the possibility that they behave similar to classic membrane lipids. However, it was suggested that the pKa of the carboxyl group of LCFAs and VLCFAs increases with the length of the acyl chain, suggesting that even at physiological pH the headgroup could be protonated^34^. However, both the protonated and the deprotonated form of VLCFAs partitioned to LDs in our simulations.

VLCFAs have been suggested to be taken up from the environment through the plasma membrane and the vectorial acylation model suggests that they need to be activated with CoA to be kept in the cell^8^. Despite previous suggestions that Fat1 transports VLCFAs through the plasma membrane and peroxisomes, its localization argues against this function, and protein modeling confirms it is not a transmembrane protein^24^. Fat1 has been identified in systematic screens as a component of LDs before^26,27,58^. We now confirm that Fat1 indeed localizes to LDs. In the absence of LDs, Fat1 localizes exclusively to the ER. Its localization depends on the N-terminal part of the protein that forms an unusually hydrophobic, yet amphipathic helix, as indicated by the observation that simple mutations altering its amphipathic nature re-localize it to the cytosol. Both *in silico* and *in vivo* experiments support a role for this motif in targeting LDs and the ER. Therefore, Fat1 and its targeting slightly deviate from the described cytosol to LDs (CYTOLD) and ER to LDs (ERTOLD) targeting pathways^45,46^, since CYTOLD-dependent proteins are described as amphipathic helix containing proteins, while ERTOLD-dependent proteins are described as containing hairpin motifs. However, several LD proteins previously described as hairpin/transmembrane domain containing proteins based on cysteine-scanning mutagenesis have recently been re-identified as monotopic integral proteins, such as the human dehydrogenase/reductase DHRS3^59^. Similar to Fat1, these proteins target LDs and the ER via a partially buried amphipathic α-helix. Other comparable targeting mechanism to that of Fat1 were previously described for the human LCFA-CoA synthetase ACSL3^60^ and one protein of the perilipin family PLIN1^61^. The similarity of Fat1 and ACSL3 targeting suggests that ACSL3 could be a direct ortholog of Fat1. However, since mammalian cells express six members each of the VLCFA CoA-synthetases, SLC27A1-A6 and the long-chain acyl-CoA synthetase (ACSL1-6), the exact homologs of Fat1 remain unclear^62^.

Our data show that the N-terminal amphipathic helix of Fat1 is not just necessary for targeting the protein to LDs or the ER, it also appears to be crucial for its enzymatic activity. Exchanging the N-terminal helix by a LD targeting motif targets Fat1 to LDs, but cannot complement its function. AF2 models predict a hydrophobic cavity that runs through the protein from the conserved active site to the membrane targeting motif. Closing this cavity by the introduction of a bulky amino acid renders the protein non-functional, even though it still localizes to LDs or the ER, respectively. We therefore propose a model **(Fig. 6)**, where the VLCFA is threaded into the hydrophobic cavity from the membrane site, similar to a VLCFA elongase from *Mycobacterium tuberculosis*^63^. This is directly dependent on the binding motif of Fat1, that creates a local membrane environment to allow the entrance of the VLCFA from the LD core. Once formed, the CoA activated product has to leave to the cytosolic opening of Fat1. Whether CoA-activated VLCFAs are soluble or if they behave similar to membrane lipids with the CoA group acting as the polar group is unknown. We have previously shown that the acyl-CoA binding protein Acb1 supports the activity of the yeast ceramide synthase, suggesting that it acts as a carrier for activated VLCFAs^64^. However, if Fat1 and Acb1 physically interact for the handover of VLCFA-CoAs remains unknown.

**Figure 6.**
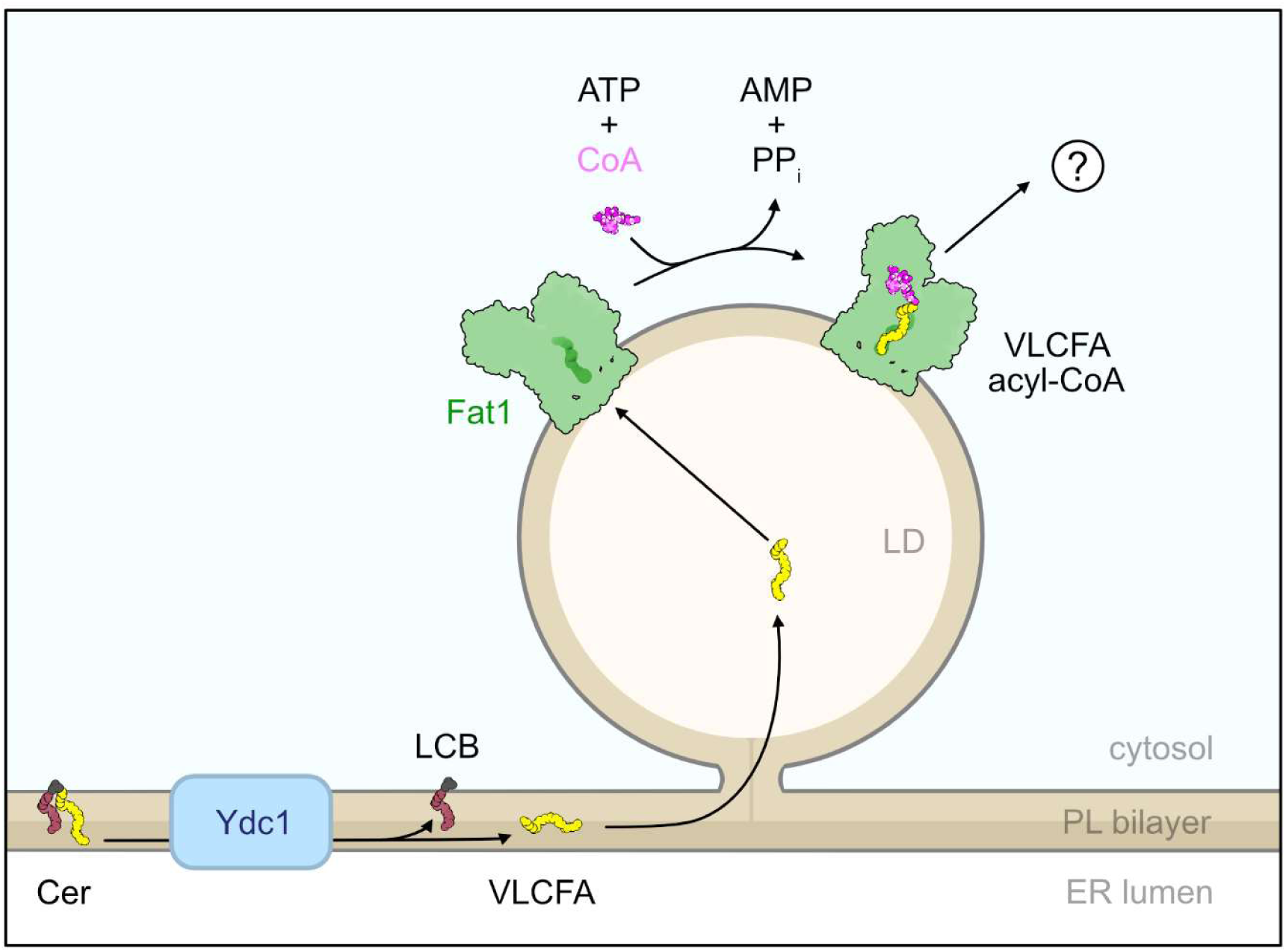
Model for VLCFA partitioning to LDs and VLCFA activation by Fat1. VLCFAs are released from ceramides by the yeast ceramidases Ydc1 (and Ypc1) at the ER membrane. Based on their high hydrophobicity, free VLCFAs partition into the hydrophobic core of LDs. Once free VLCFAs encounter the hydrophobic cavity of Fat1, including its amphipathic helix at the monolayer interface, VLCFAs can enter the active site to be activated with CoA in an ATP-dependent manner. How the activated VLCFA acyl-CoA product is released from Fat1 and how it reaches its cellular destinations remains unclear.

Although Fat1’s enzymatic function as a VLCFA-CoA synthetase is well-established, the cellular impact of its function remains elusive. Our genetic analyses demonstrate that Fat1 is necessary to maintain sufficient VLCFA levels when *de novo* VLCFA biosynthesis is impaired by deleting the FA elongase Elo2. A recent study suggested that Fat1 is required to avoid membrane saturation and induction of the unfolded protein response (UPR)^65^. This could be particularly important under conditions where cells lack LDs. Accumulation of free VLCFAs could cause membrane stress and induction of the UPR. Interestingly, mis-regulation of the VLCFA elongase Elo2 results in increased synthesis of VLCFAs. These CoA-activated, over-produced VLCFAs are also incorporated into TGs^5^. Consequently, Fat1 function would also be essential when VLCFAs are released from TGs. Deletion of Fat1 results in VLCFA accumulation suggesting that LD-localized Fat1 may activate free VLCFAs generated by TG hydrolysis^25,32^. Substrate shuttling by FA-CoA synthetases at sites of increased lipid biosynthesis has also been proposed^58^.

Finally, all the components described in our model are conserved throughout evolution. Mammalian cells express LCFA elongases that also localize to LDs. Similarly, the mammalian acyl-CoA-binding protein has also been suggested to be required for optimal ceramide synthase activity^66^. In humans, VLCFAs have been implicated in diseases such as X-linked adrenoleukodystrophy^67–69^. That the targeting mechanism of Fat1 or other CoA-synthetases has co-evolved with the biophysical properties of VLCFAs, together with a better understanding of the exact mechanism of VLCFA activation, will help us to understand the role of VLCFAs in both cell physiology and physiopathology.

## Methods

### Yeast strains and plasmids

Yeast strains, plasmids, and oligonucleotides used in this study are listed in Supplementary Tables 1, 2, and 3.

### Genetic interactions

To conduct tetrad analyses, diploid yeast cells were collected by centrifugation and placed onto 1% potassium acetate agar plates for sporulation at 30°C. After 3-5 days and microscopic inspection for ascus formation, a sample of each culture was suspended in 100 µl of sterile water. 5 µl of Zymolyase 20T (10 mg/mL; MP Biomedicals, Eschwege, Germany) were added, and incubated at room temperature for 9 minutes. A small amount of cells was streaked out on YPD plates and spores were segregated using a Singer MSM400 micromanipulator (Singer Instruments, Somerset, UK). The plates were then incubated for 3 days at 30°C.

### Fluorescence microscopy

Cells were grown overnight at 30 °C in synthetic medium supplemented with essential amino acids (SDC) and grown from an OD_600_ of 0.2 to logarithmic growth phase the next morning. Cells were centrifuged at 4,000 rpm for 5 mins, resuspended in fresh SDC medium and directly imaged live on an Olympus IX-71 inverted microscope equipped with a 100x numerical aperture (NA) 1.49 oil-immersion objective, a sCMOS camera (PCO, Kelheim, Germany) and an InsightSSI illumination system. Data were acquired with SoftWoRx software (Applied Precision, Issaquah, WA) followed by constrained-iterative deconvolution (SoftWoRx) and processed with ImageJ 2.1.0. (National Institutes of Health, Bethesda, MD; RRID:SCR_003070).

### MD setups and simulations

Droplet-in-bilayer systems were prepared starting from AA-bilayers with different compositions: DOPC 95% - VLCFAs charged 5%; DOPC 95% - VLCFAs neutral 5%; DOPC 95% - VLCFAs charged 2.5% - VLCFAs neutral 2.5%. The bilayers were generated and equilibrated using CHARMM - graphical user interface (GUI) *membrane builder* tool^70,71^, then the two leaflets were separated using TopoTools^72,73^, and finally replicated in order to reach the desired dimensions (∼57×57 nm). In between the two monolayers, a box containing 1440 AA-TO lipids, previously minimized and equilibrated using CHARMM36m force field^74–76^ and the GROMACS software^77^, was placed. This configuration was mapped to CG-SPICA using the *cg spica tool* (https://zenodo.org/records/10611579) and equilibrated for 12 ns using LAMMPS software and employing the SDK/SPICA force field^40,78,79^. Parameters for VLCFAs were adapted from MacDermaid et al 2020^80^, while parameters for the TO molecules were taken from Campomanes et al, 2019^81^. Finally, two replicates for each system were run for 800 ns using a timestep of 20 fs. During the production, the temperature was kept at 310 K using a Nosé-Hoover thermostat^82^, while the pressure control was maintained at 1 atm employing the Nosé-Hoover barostat^83^. Linear and angular momenta were removed every timestep. Van der Waals and electrostatic interaction were truncated at 1.5 nm; long-range electrostatics beyond the same cutoff were computed using the particle-particle-particle-mesh solver, with a root mean square force error of 10^−5^ kcal mol^−1^ Å^−1^ and order 3.

The droplet-in-bilayer system enriched with VLCFAs charged 2.5% - VLCFAs neutral 2.5% was also simulated in the presence of the yeast Fat1. The protein, modeled based on its AF2 prediction (uniprot accession number: P38225), was initially positioned in solution with a random orientation and at nine different starting point. Duplicates of each configuration of were run for 1.2 μs with a timestep of 10 fs, using the same conditions described above.

Fat1 was also simulated in presence of a pure DOPC bilayer and trilayer composed by DOPC monolayers and pure TO oil core. The system containing Fat1 in presence of a simple DOPC bilayer was initially generated using the CHARMM-GUI^70,71^ *membrane builder* module, with the protein being randomly placed in solution at around 6 nm distance from the membrane. The system was later re-solvated and ionized with a 0.15 NaCL solution to get rid of excess water. Upon equilibration following the default CHARMM-GUI 6-step protocol, the atomistic system was converted to SPICA-CG model and subsequently further minimized and equilibrated. Three replicates were simulated for a total of about 7 μs, using a timestep of 10 fs. Trilayer systems in presence of Fat1 were prepared as follow: first, a small (3×3 nm) pure DOPC-AA bilayer was generated using CHARMM-GUI *membrane builder*, then minimized and equilibrated using the classical CHARMM-GUI workflow. Once equilibrated, the bilayer was first replicated and then the two leaflets were separated. Later, a previously assembled and equilibrated box with containing 718 AA-TO molecules was placed between the two separated monolayers. The protein was placed in solution at around 4 nm distance from the monolayer surface and a cycle of minimization and equilibration was run. The equilibrated system was then mapped into CG-SPICA, and three replicas were first equilibrated and then simulated for 2 μs of production each with a timestep *d*t=10 fs.

MD productions for both bilayer and trilayer in presence of Fat1 were run under the same conditions described above for the droplet-in-system.

### Analysis

Cavity pocket detection was performed utilizing the Fpocket software, an algorithm relying on Voronoi tessellation^84^, giving as a reference structure for the pocket identification the AF2 model of Fat1. The same model has been used for the analysis of the evolutionary conservation employing the Consurf tool^85^.

The radial distribution of VLCFAs (neutral and anionic) was performed using MD Analysis^86,87^ by measuring the distance of their headgroup beads from the blister TO center of mass (COM) overtime. The distributions were generated using the average per-frame distances of both the lipid species. The DOPC headgroups were utilized to define the average radius of the phospholipids shell encircling the TO core.

The protein-membrane association for the trilayer system was calculated using the center of mass of the protein and the membrane, and the distance between them was measured using the GROMACS *gmx pairdist* module. The occupancy map to identify the protein-membrane interface was generated using *gmx select*, using a trajectory concatenated over 3 replicas.

For the ternary system, the distance between the centers of geometry (COG) of the protein and the TO-core was calculated over time for each of the 18 replicas, considering only the x and y coordinates. Each 1.2 μs simulation was divided into six equal chunks (each corresponding to 0.2 μs), and distances obtained from all independent simulation replicas were combined within each time window.

Enrichment-depletion analysis on ternary systems (both in the absence and in the presence of the Fat1 protein) was conducted as in Jimenez-Rojo et al., 2020^88^. 2D lateral density maps were first generated for each lipid component using *gmx densmap*. Then, each density was divided by the sum of the densities of the other membrane species and finally divided by the concentration of each lipid component.

Over time protein x/y-tracing were computed aligning the protein COM trajectories to the droplet COM and projecting them onto the x/y plane, with the time normalized from start to end. Per each replica, only the first 1 μs after the protein was bound to the ternary system surface was considered.

Fat1 structure prediction in presence of a VLCFA ligand were run utilizing Alphafold3 (AF3)^50^ and Boltz-2 (Rowan Scientific. https:/www.rowansci.com)^51^.

All the molecular images were rendered using Visual Molecular Dynamics (VMD) software^73^ and all plots were generated using the python module matplotlib^89^.

### Fluorescence spectroscopy of neutral lipid emulsions

Neutral lipid emulsions were prepared by mixing 15 μl of neutral lipids with 300 μl of buffer. Three lipid compositions were tested: (i) pure triacylglycerol (TAG; G7793, Sigma-Aldrich), (ii) TAG supplemented with oleic acid (OA; O1383, Sigma-Aldrich) at 95/5% (w/w), and (iii) TAG supplemented with hexacosanoic acid (HCA) at 95/5% (w/w). Two buffer conditions were used: either a 0.1 mol·L⁻¹ NaOH solution or a HKM buffer composed of 50 mM HEPES, 120 mM potassium acetate, and 1 mM MgCl₂ prepared in Milli-Q water and adjusted to pH 7.4 and 270 mOsm. Emulsification was performed using a bath sonicator until a visually homogeneous emulsion was obtained. The emulsions were then transferred to a microplate and analyzed using a spectrofluorometer. Excitation spectra were recorded between 210 and 350 nm, and a fluorescence emission peak was consistently detected at 340 nm.

### Protein purifications

Purification of 3xFLAG tagged Fat1 was performed as previously described^3^. Yeast cells were harvested after growth for 24 h in a yeast peptone (YP) medium containing 2% galactose (v/v). The harvested cells were resuspended in lysis buffer (50 mM HEPES-KOH (pH 6.8), 150 mM KOAc, 2 mM MgOAc, 1 mM CaCl_2_, 200 mM Sorbitol). supplemented with 1mM phenylmethylsulfonylfluoride (PMSF) and 1x FY protease inhibitor mix (Serva) in a 1:1 ratio (w/v) and afterwards frozen dropwise in liquid nitrogen. The frozen cells were lysed using a 6875D Freezer/Mill Dual-Chamber Cryogenic Grinder (SPEX SamplePrep) for 15 cycles of 2 mins at 12 CPS. The resulting yeast powder was thawed in lysis buffer containing 1mM PMSF and 1x FY. Cell debris was removed by two centrifugation steps at 1,000 *g* at 4 °C for 20 mins. Microsomal membranes were collected at 44,000 *g* at 4 °C for 30 mins, resuspended in lysis buffer and then diluted with IP buffer (50 mM HEPES-KOH (pH 6.8), 150 mM KOAc, 2 mM MgOAc, 1 mM CaCl2, 15 % Glycerol) supplemented with 1% glyco-diosgenin (GDN) and protease inhibitors. After solubilization for 1.5 hours at 4 °C and a centrifugation step at 44,000 *g* at 4 °C for 30 mins, the supernatant was added to α-FLAG resin (Sigma Aldrich). Beads were washed twice with IP buffer containing 0.1 % and 0.01 % GDN, respectively, after incubation for 45 mins at 4 °C. 3xFLAG peptide was added twice and incubated together with α-FLAG beads for 45 mins and 5 mins on a turning wheel. Proteins were eluted at 460 *g* at 4 °C for 30 s.

### Fat1 *in vitro* activity assay

Fat1 activity was measured by detecting AMP levels released during the Fat1-catalyzed formation of a thioester between CoA and fatty acid. The assays were performed in a MS glass vial in a total volume of 100 µl. The assay consists of IP buffer, 0.008 % GDN, 300 µM DTT, 2 mM MgCl_2,_ 0.01% Triton, 100 µM ATP, 150 mM CoA and 10 µM fatty acid substrate. The fatty acid (C26:0 and C16:0, Cayman Chemical) substrates were dried in a Speedvac at 45 °C, resuspended in 200µl 50 mM HEPES-NaOH (pH 6.8) containing 10 mg/ml α-cyclodextrin (Carl Roth) and incubated for 30 min in a sonicating water bath at room temperature. The enzymatic reaction was initiated by adding purified protein. After incubation at 26°C for 30 min, the samples were deproteinized with a 10 kDa MWCO concentrator (Merck Millipore). To measure AMP levels, 25 µl of each sample was transferred to a 96-well plate and AMP levels were determined using the AMP-Glo Assay Kit (Promega). The resulting luminescence signal was measured in a SpectraMax iD3 Multi-Mode microplate reader with 140 ms integration time. Background luminescence signal of protein-free samples were subtracted from the measurements of protein-containing samples.

### Mass spectrometry-based expression test

Yeast cells were grown in 10 ml YPD medium at 30°C until they reached the logarithmic growth phase. One OD unit of each cell culture was harvested via centrifugation for 5 min at 4,000 g. Cell lysis was performed in 50 µl lysis buffer of the PreOmics iST kit by placing the samples in a heating block at 95 °C, 1000 rpm for 10 min. The lysates were digested by incubation O/N at 37 °C at 500 rpm with Trypsin/LysC and further prepared following the instructions of the iST Sample Preparation Kit (PreOmics). The dried peptides were resuspended in 500 μl LC-Load and 2 µl of each peptide sample were used for analysis by reversed-phase chromatography as previously described^4^ using a Thermo UltiMate 3000 RSLCnano system coupled to a TimsTOF HT mass spectrometer (Bruker Corporation, Bremen, Germany) via a Captive Spray Ion source. Peptide separation was achieved on an Aurora Gen3 C18 column (25 cm × 75 µm, 1.6 µm) with CSI emitter (Ionoptics, Australia) at 40 °C. Elution was performed using a linear gradient of acetonitrile from 2 - 35 % in 0.1 % formic acid over 44 min at a constant flow rate of 300 nl/min, followed by an increase to 50 % over 7 min and to 85 % Buffer B over 4 min. The eluted peptides were directly electro sprayed into the mass spectrometer at an electrospray voltage of 1.5 kV and 3 l/min Dry Gas. The TimsTOF HT settings were adjusted to positive Ion polarity with a MS range from 100 to 1700 m/z. The scan mode was set to DDA-PASEF. 10 PASEF ramps per cycle resulted in a duty cycle time of 1.17s. The target intensity was adjusted to 14,000. The intensity threshold was set to 1,200. The dynamic exclusion time was set to 0.4 min. Precursor Ion charge state was limited from 0 to 5. For MS data analysis the resulting data were processed using MaxQuant (V2.7.0.0, www.maxquant.org) and subsequent statistical analyses were performed with Perseus (V2.1.5.0, www.maxquant.org/perseus)^90–92^. Significance thresholds in the volcano plot were determined by a permutation-based false discovery rate (FDR) method. The mass spectrometry proteomics data have been deposited to the ProteomeXchange Consortium via the PRIDE partner repository with the dataset identifier PXD070847 and 10.6019/PXD070847^93,94^.

### Statistical analyses

Colocalization analysis was performed on microscopy images subsequent to their constrained-iterative deconvolution (SoftWoRx) using the BIOP JACoP (Just Another Colocalization Plugin) plugin^95^ implemented in Fiji^96^. Regions of interest (ROIs) were manually defined to include only dividing cells. Three independent datasets were analyzed comprising a total of 100 cells. The Pearson correlation coefficients obtained for individual ROIs were exported and the mean value for each data set was calculated using Excel (2019). Data from *in vitro* activity assays were analyzed using Microsoft Excel (2019). For each dataset, the mean and standard deviation (SD) were calculated. Results are presented as mean ± SD.

## Acknowledgments

We thank members of the F.F. and S.V. labs for their valuable comments on the manuscript. S.V. acknowledges support by the Swiss National Science Foundation (310030_219264). This work was supported by grants from the Swiss National Supercomputing Centre (CSCS) under project ID s1176 and lp24 to S.V. F.F. is supported by the German Research Foundation (DFG); project numbers 491484150, 503478512, 467522186, 467522186 and the *RISE UP!* program of the Boehringer Ingelheim Foundation.

## Supplementary material

**Table 1:**
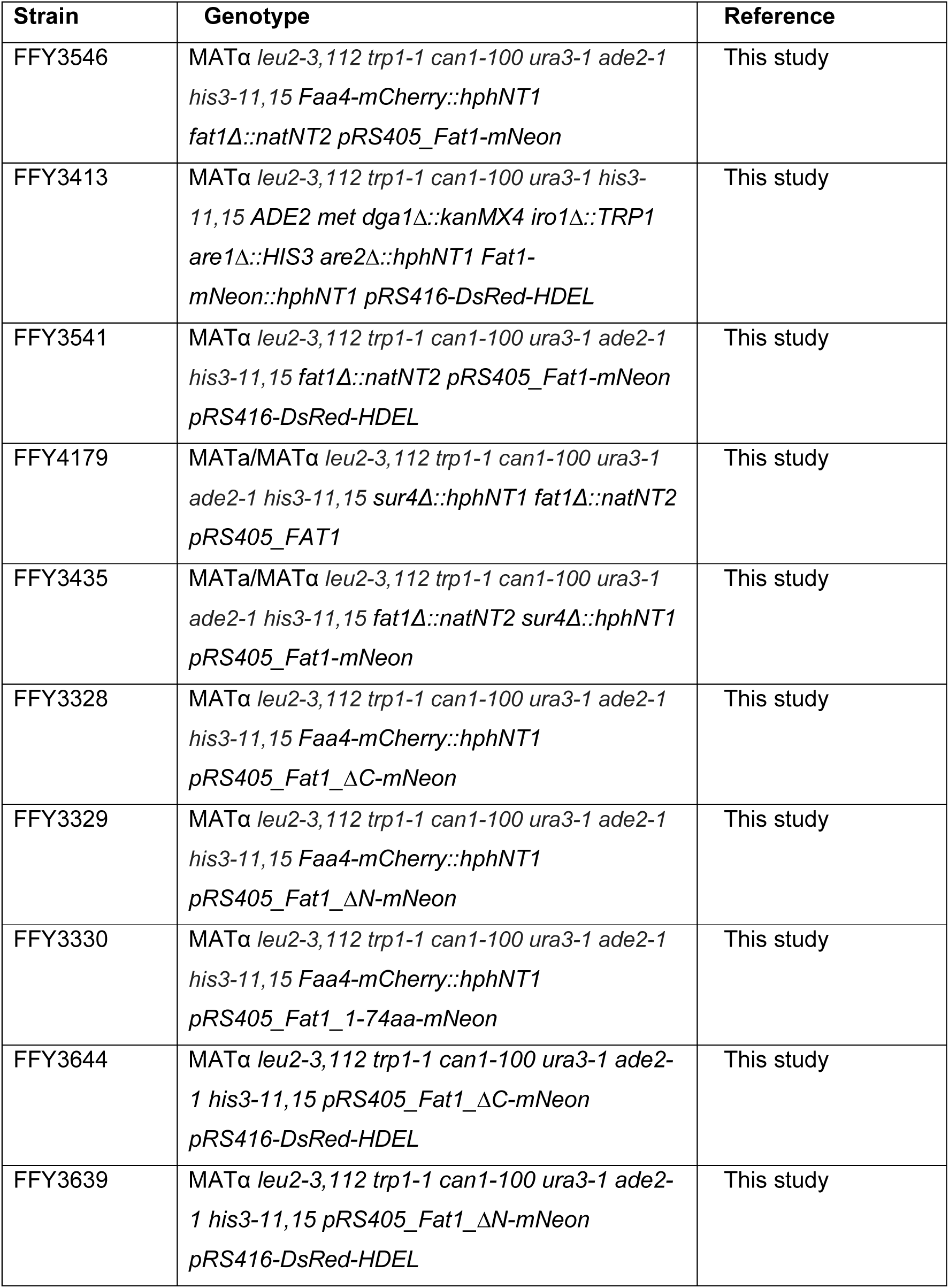
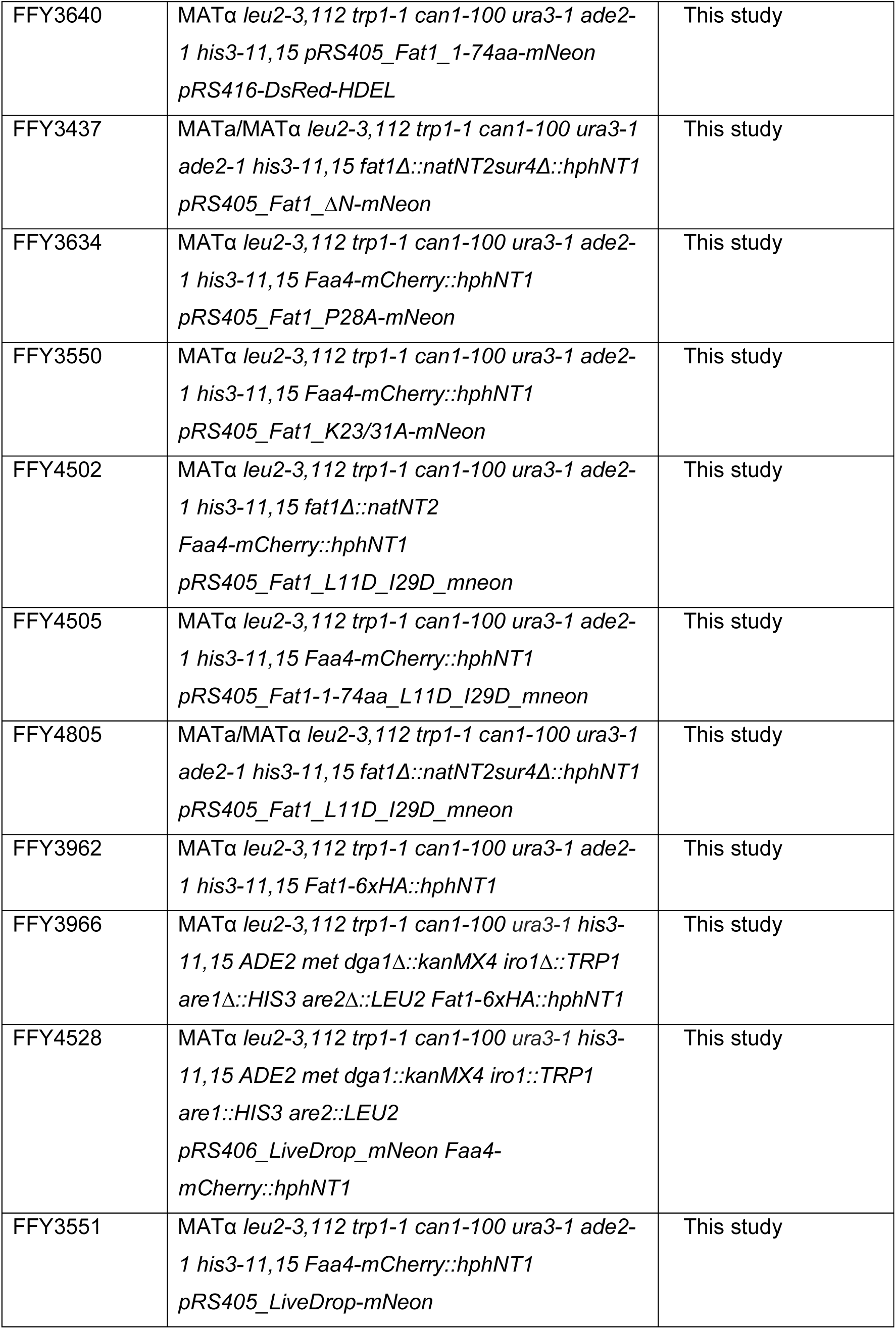
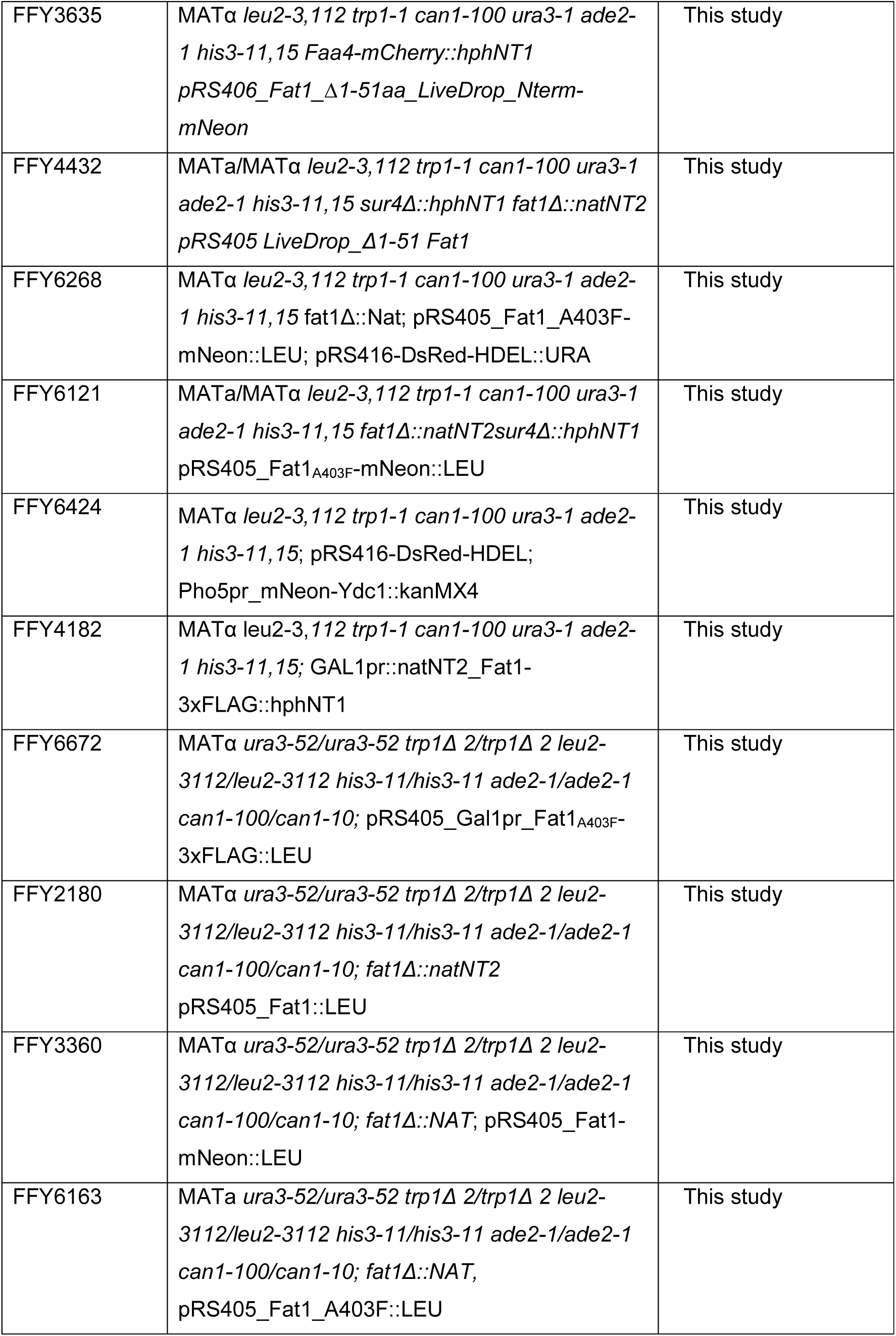
List of all yeast strains and their genotypes used in this study.

**Table 2:**
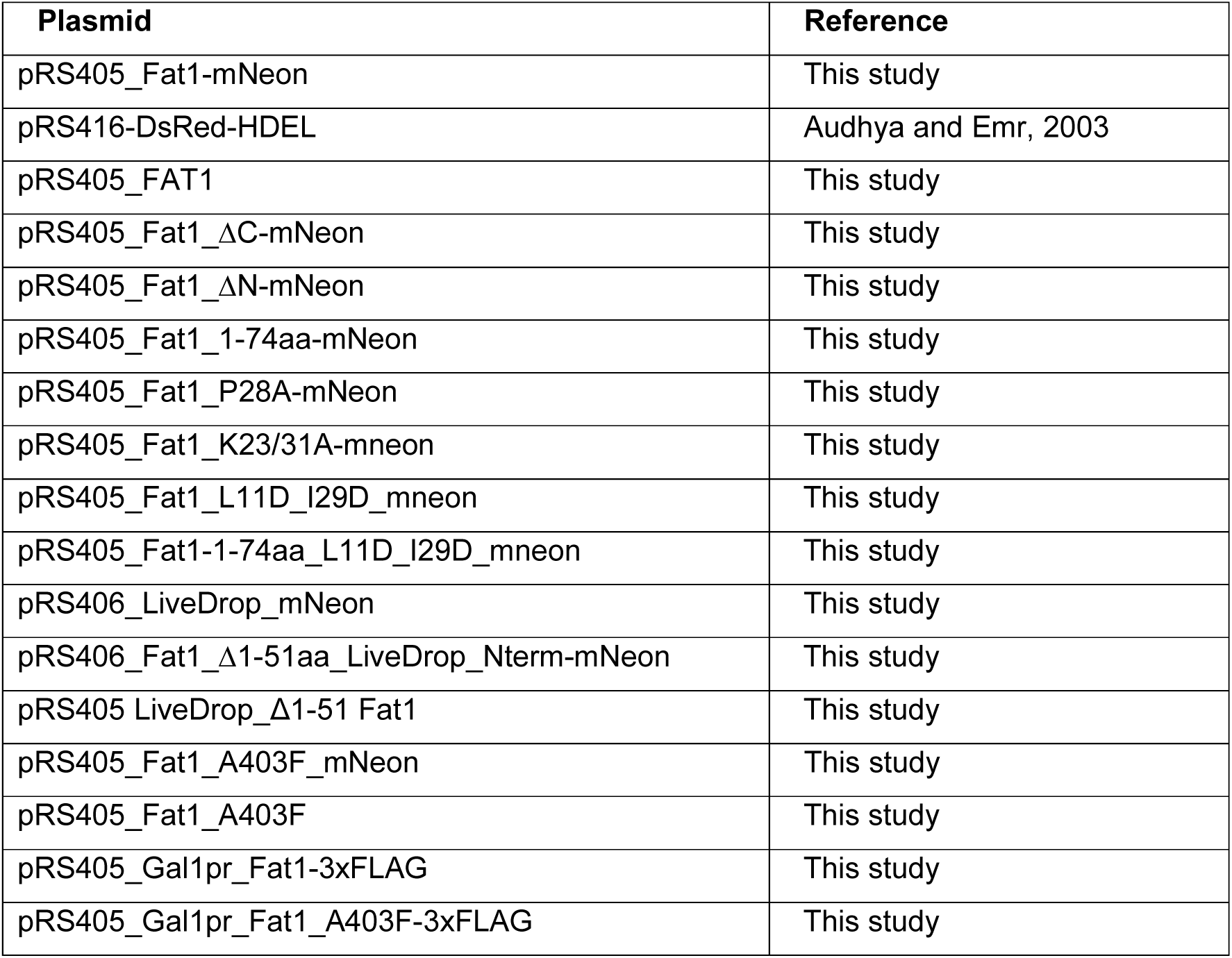
List of all plasmids used in this study.

**Table 3:**
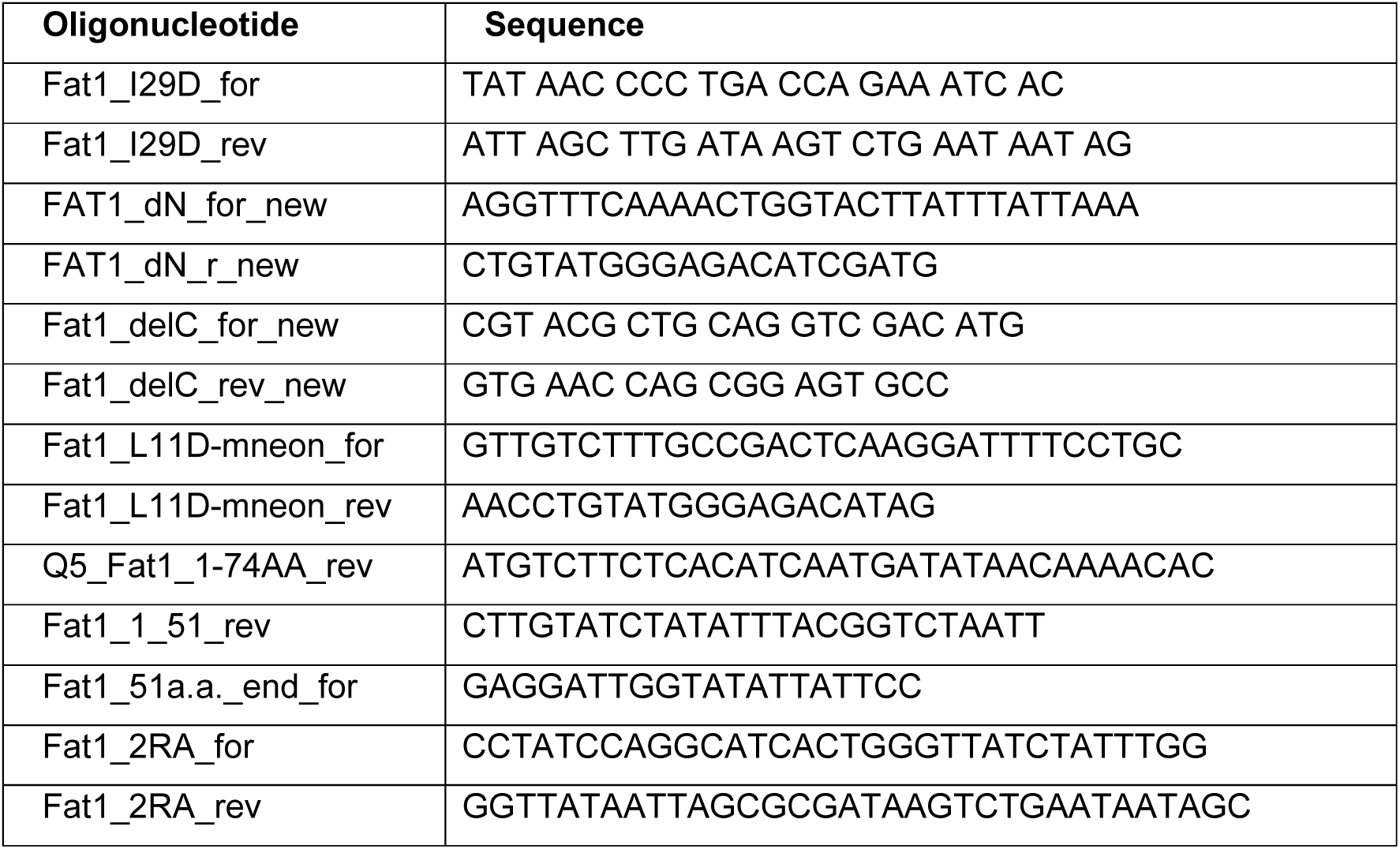
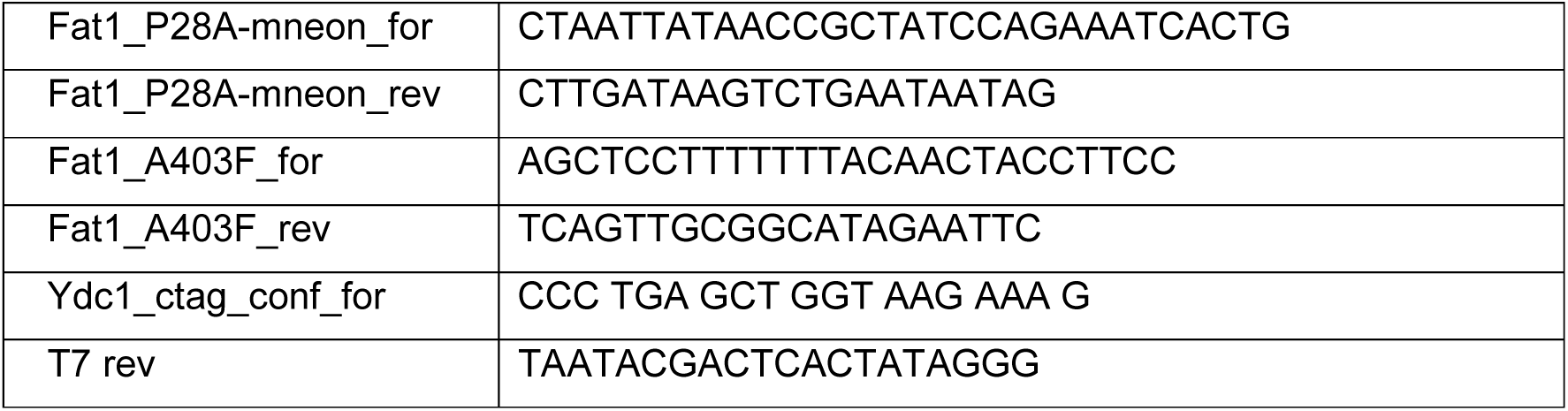
List of all oligonucleotides used in this study.

**Table 4:** Raw Data Proteomics

## Supplementary Figures

**Supplementary Figure 1.**
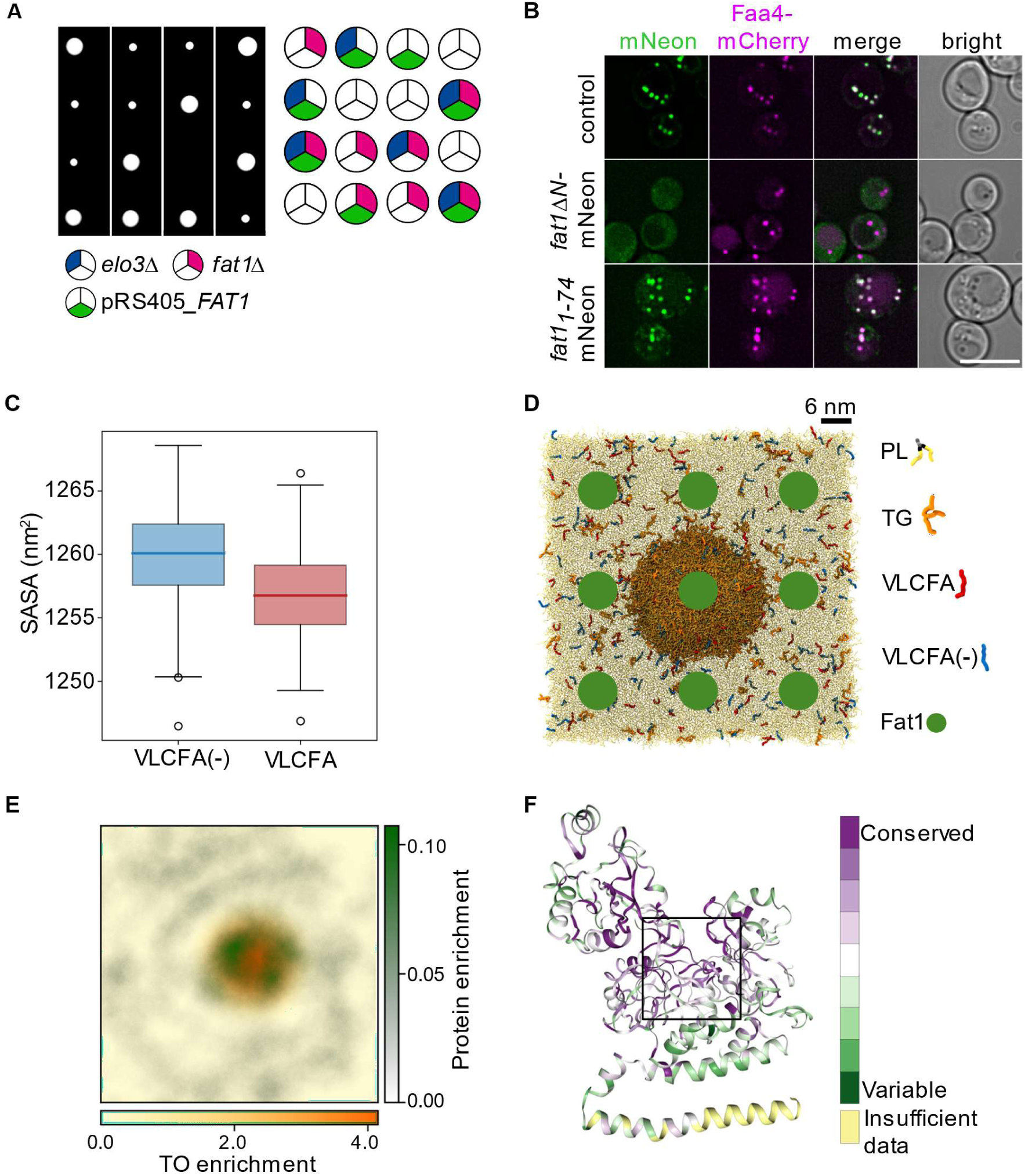
**A)** Tetrad analysis of *fat1Δ* pRS405-*FAT1* cells (pink and green, respectively) mutants crossed with *elo3Δ* (blue). **B)** Co-localization of mNeon-tagged WT Fat1 (upper panel), Fat1-mNeon lacking the first 74 amino acids (*fat1ΔN*; middle panel) and the 74 Fat1-N-terminal amino acids fused to mNeon (*fat1_1-74_*-mNeon, lower panel) with Faa4-mCherry. Representative confocal midsections are shown (upper panel). Scale bar = 5 µm. **C)** SASA analysis shows an increased exposition of VLCFAs towards the water solvent in an ER-LD like system. **D)** ER-LD like system’s top view showing the 9 configurations of Fat1 (green dot) initially positioned in solution. Two replicates of such 9 configurations have been simulated. **E)** Protein number density analysis shows the binding preference of Fat1 (green) towards the monolayer (marked by the presence of a TO blister, orange) over bilayer. **F)** Consurf ^85^ prediction of the Fat1 binding site.

**Supplementary Figure 2.**
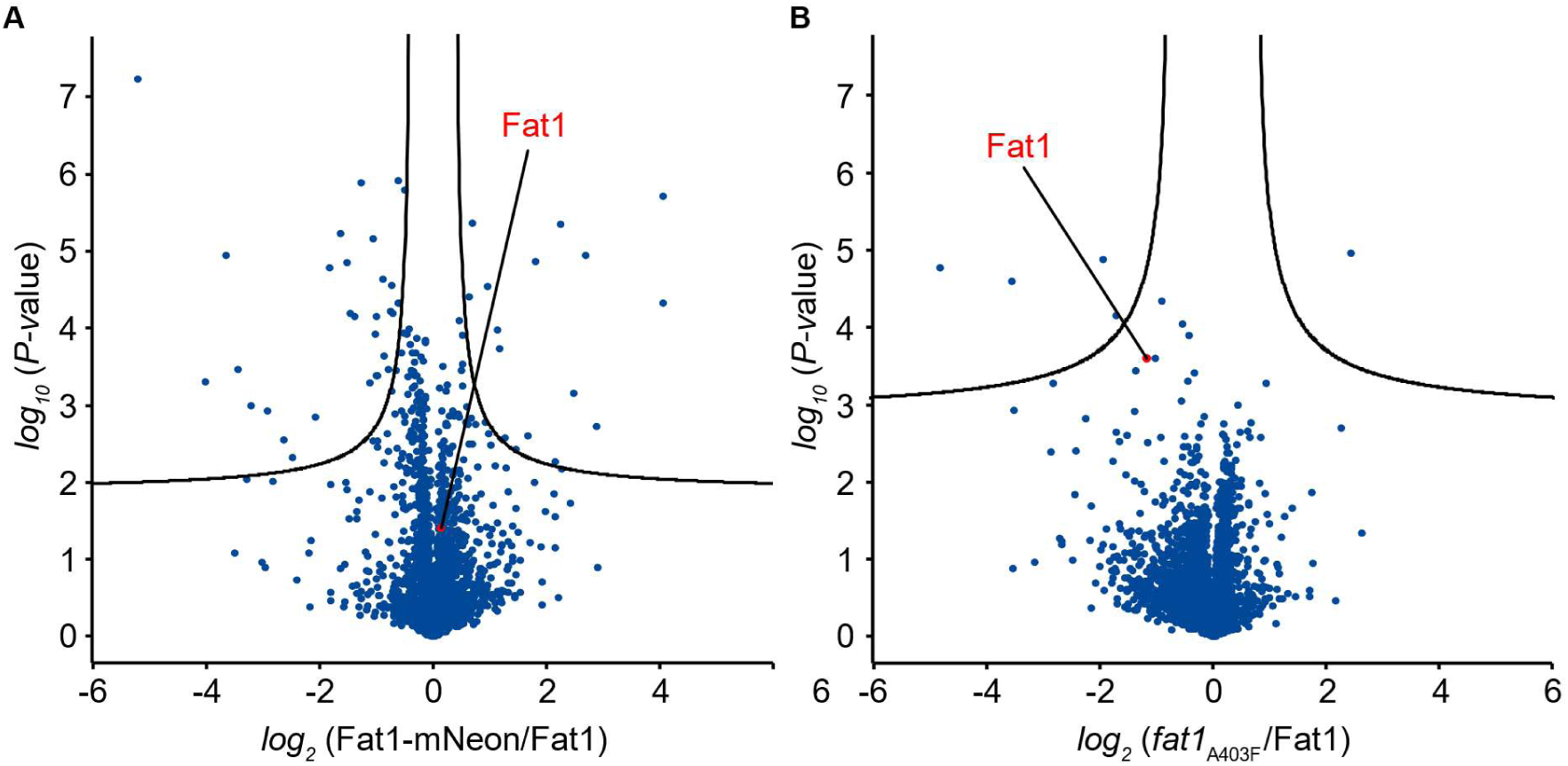
**A)** Label free proteomics of yeast cells expressing plasmid-introduced Fat1-mNeon compared to yeast cells expressing plasmid-introduced WT Fat1. In the volcano plot, the protein abundance ratios of Fat1-mNeon over control cells are plotted against the negative *log*_10_ of the *P*-value of the two-tailed t-test for each protein. **B)** Label free proteomics of yeast cells expressing plasmid-introduced *fat1*_A403F_ compared to yeast cells expressing plasmid-introduced WT Fat1. In the volcano plot, the protein abundance ratios of *fat1*_A403F_ over control cells are plotted against the negative *log*_10_ of the *P*-value of the two-tailed t-test for each protein.

